# 3DCellComposer - A Versatile Pipeline Utilizing 2D Cell Segmentation Methods for 3D Cell Segmentation

**DOI:** 10.1101/2024.03.08.584082

**Authors:** Haoran Chen, Robert F. Murphy

**Affiliations:** Computational Biology Department, School of Computer Science, Carnegie Mellon University, 5000 Forbes Avenue, Pittsburgh PA 15213, USA

**Keywords:** cell segmentation, nuclear segmentation, spatial proteomics, multiplexed fluorescence imaging, imaging mass spectroscopy, 3D microscopy, tissue images

## Abstract

**Background:** Cell segmentation is crucial in bioimage informatics, as its accuracy directly impacts conclusions drawn from cellular analyses. While many approaches to 2D cell segmentation have been described, 3D cell segmentation has received much less attention. 3D segmentation faces significant challenges, including limited training data availability due to the difficulty of the task for human annotators, and inherent three-dimensional complexity. As a result, existing 3D cell segmentation methods often lack broad applicability across different imaging modalities.

**Results:** To address this, we developed a generalizable approach for using 2D cell segmentation methods to produce accurate 3D cell segmentations. We implemented this approach in 3DCellComposer, a versatile, open-source package that allows users to choose any existing 2D segmentation model appropriate for their tissue or cell type(s) without requiring any additional training. Importantly, we have enhanced our open source CellSegmentationEvaluator quality evaluation tool to support 3D images. It provides metrics that allow selection of the best approach for a given imaging source and modality, without the need for human annotations to assess performance. Using these metrics, we demonstrated that our approach produced high-quality 3D segmentations of tissue images, and that it could outperform an existing 3D segmentation method on the cell culture images with which it was trained.

**Conclusions:** 3DCellComposer, when paired with well-trained 2D segmentation models, provides an important alternative to acquiring human-annotated 3D images for new sample types or imaging modalities and then training 3D segmentation models using them. It is expected to be of significant value for large scale projects such as the Human BioMolecular Atlas Program.

## Background

The development of multiplexed imaging methods enables biomedical studies at the cellular level from numerous perspectives using various markers that target specific proteins [1-4]. The recent emergence of 3D multiplexed tissue imaging techniques further provides unprecedented potential for investigating the interaction, organization, and specialization of cells in 3D tissues [5, 6]. However, realizing that potential also requires accurate 3D cell segmentation methods in order to ensure the quality of subsequent cellular analyses.

Within the challenging domain of 3D cell segmentation, tasks related to nuclear segmentation have received more attention. The main reason is that 3D nuclei segmentation is easier than 3D cell segmentation, primarily because of the well-defined round shape of nuclei and the high clarity provided by nuclear dyes. Over the years, traditional algorithms such as watershed [7], wavelet [8], gradient flow tracking [9], and a multi-model approach combining more than one traditional algorithm [10] have been employed for efficient 3D nuclei segmentation. Recently, deep learning-based methods have been employed to segment 3D nuclei more accurately with specifically tailored neural network architectures [11-14]. While 3D nuclei segmentation may be useful by itself in some applications, 3D cell segmentation is clearly needed in many others.

However, 3D cell segmentation comes with its own set of challenges. One major issue is that the shapes and boundaries of whole cells are often less well-defined than those of nuclei. This problem is particularly pronounced in tissue images, where the consistency and quality of cell membrane or cytoplasmic markers can vary significantly between different samples and imaging modalities. Various studies have explored approaches to tackle 3D cell segmentation, although not all focus on multiplexed imaging. Eschweiler et al. [15] explored the use of the spherical harmonics technique as an alternative approach to shape-constrain the star-convex segmentation prediction of 3D microscopy image data. Wolny et al. [16] introduced Plantseg, which combines a 3D U-Net with region adjacency graph partitioning to predict cell boundaries in 3D plant tissue images. Wang et al. [17] developed 3DCellSeg, which employs a lightweight U-Net-like network with a custom loss function, aiming to predict and de-clump 3D cells. The Allen Cell and Structure Segmenter (referred to below as ACSS) presented a human-in-the-loop deep learning workflow with expert correction of the segmentation results at each iteration [18, 19]. One significant limitation common to these methods is their reliance on deep neural networks to handle the complexity of three-dimensional tissue data. Such approaches require large amounts of training data that might not always be available, especially when a new 3D imaging modality emerges. Even if certain 3D imaging modalities may offer sufficient images for deep learning training, their variety and volume are still limited compared to the extensive libraries of 2D cell images. This limitation could reduce the applicability of the 3D cell segmentation model, meaning that it might perform well on the 3D image modality it was trained on but not on other modalities. Furthermore, training these models demands extensive manual labeling of 3D cells by experts [20], which inevitably suffer from intra and inter-expert variability [21, 22]. Human-in-the-loop methods are designed to reduce the need of manual labeling, but still demand a considerable amount of expert input [18].

On the other hand, 2D cell segmentation models, for both nuclei and cell segmentation, are often trained with datasets (e.g., TissueNet [23] and LIVECell [24]) containing millions of cells in 2D tissue and cell culture images across multiple imaging modalities. This allows for an alternative way to approach the 3D cell segmentation problem by using a 2D model to segment images and then assembling the 3D cells. One variation on this approach has been incorporated into Cellpose [25, 26], which applies its 2D segmentation capability to individual slices along three axes and then averaging the resulting spatial gradients to derive 3D cells. However, this approach is tailored specifically to the spatial gradient technique used by Cellpose, limiting its use with other algorithms and reducing its applicability to image types that are not part of Cellpose’s original training data.

To overcome these challenges, we introduce 3DCellComposer, a versatile tool capable of utilizing any 2D cell segmentation model for 3D cell segmentation tasks. Instead of retraining 3D deep learning cell segmentation models using a limited number of 3D images, 3DCellComposer takes advantage of well-trained 2D models without requiring any further training. This allows users to customize their analysis pipelines by selecting a 2D segmentation model extensively trained from a similar tissue or imaging modality as their 3D data, or any other model that suits their needs. To further assist users in selecting the most effective 2D segmentation model for their 3D images, we have enhanced our earlier segmentation quality evaluation tool, CellSegmentationEvaluator [27] to support 3D segmentations. This combination provides a single solution for 3D cell segmentation needs. Figure 1 illustrates the whole pipeline with specific steps involved in segmenting 3D cells using our software.

**Figure 1.**
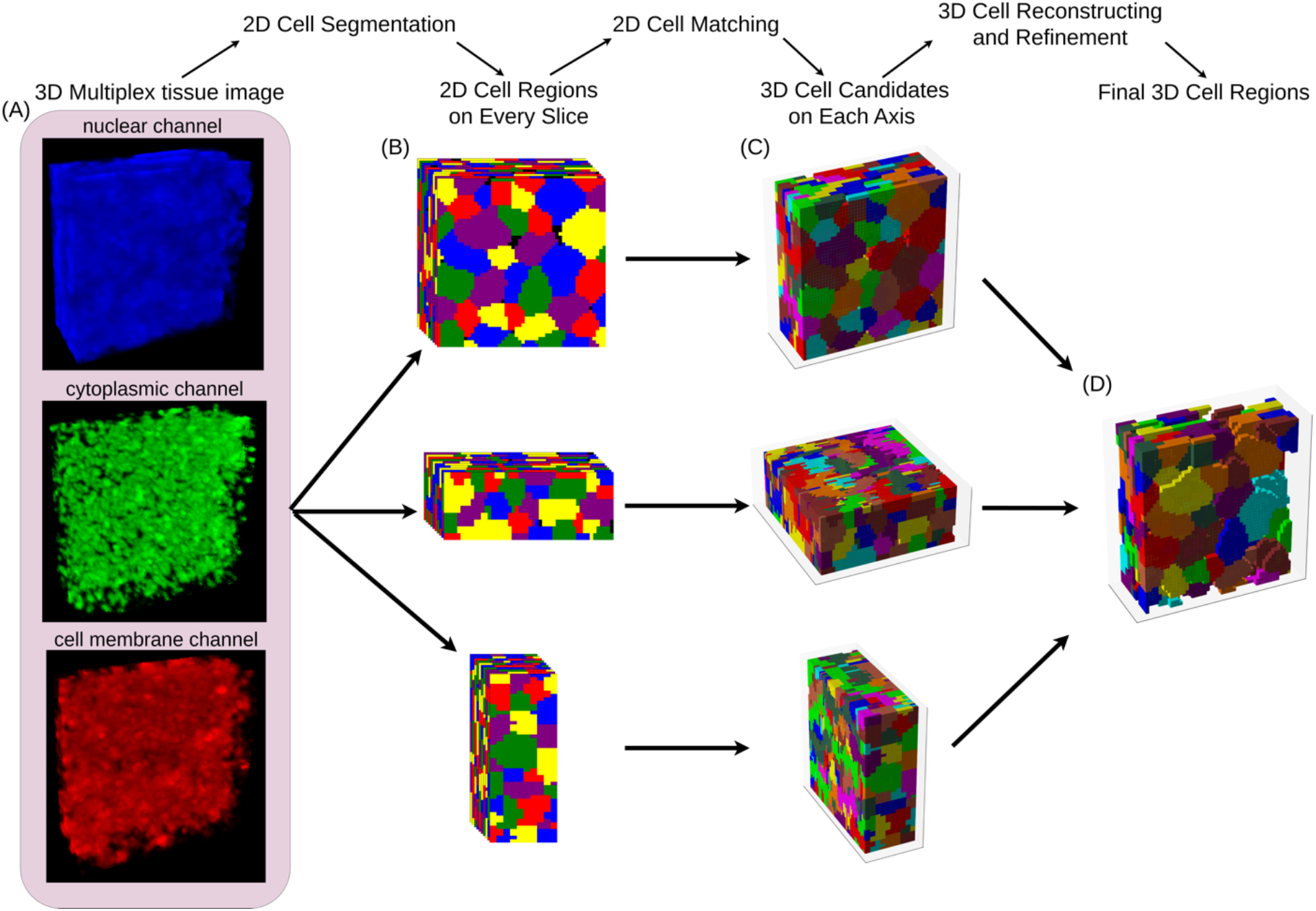
Overview of the 3DCellComposer pipeline for 3D cell segmentation. (A) A 3D multiplexed image serves as the input for segmentation. The image channels for the nucleus, cytoplasm, and cell membrane are shown in different colors. (B) A 2D cell segmentation method of user’s choice is applied to every slice along all three axes. Adjacent cells are colored differently for distinction. (C) 2D cell matching is conducted on each axis to generate 3D cell candidates. (D) The final 3D cell regions are obtained by performing linear indexing reconstruction and subsequent refinement.

We tested our pipeline using 6 distinct 2D cell segmentation methods, covering both traditional and deep learning-based techniques, and benchmarked their performance against previously-described 3D cell segmentation methods, including Cellpose 2D to 3D, ACSS, and 3DCellSeg (Supplementary Table 1). Using the enhanced CellSegmentationEvaluator, we computed a segmentation quality score for each method to objectively assess its performance without costly manual annotations for 3D cells. Our analyses indicate that 3DCellComposer with a well-trained 2D segmentation model can yield accurate segmentation of 3D cell boundaries across diverse tissues in 3D multichannel tissue images. In contrast, existing 3D segmentation methods often failed to segment cells or excessively merged cells from these images. Furthermore, 3DCellComposer also showed competitive performance against ACSS when applied to cell culture images from the same cell line and modality as ACSS’s own training data.

## Methods

### 3D multiplexed tissue and cell culture images

We obtained three 3D Imaging Mass Cytometry (IMC) [28, 29] datasets from the Human BioMolecular Atlas Program (HuBMAP) database [30, 31], corresponding to spleen, thymus, and lymph node tissues, respectively (see Supplementary Table 2 for details). These images were produced by the University of Florida Tissue Mapping Center.

For the segmentation process, we selected Iridium as the nuclear channel. The cytoplasmic channel was generated by summing the signals from three markers: Smooth Muscle Actin (SMA), Granzyme B, and Myeloperoxidase (MPO). For the cell membrane channel, we created a composite by summing the signals from five different markers: E-Cadherin/P-Cadherin, pan-Cytokeratin, CD31, CD45RA, and CD45RO (see Supplementary Table 3 for details). We chose to use multiple markers for the cytoplasmic and cell membrane channels because there are no universal markers that target these two cellular components across all cells provided in IMC data. By selecting and combining a diverse range of markers, each targeting one or multiple cell types, we aimed to cover as many cells as possible within the tissue for cell segmentation. Notably, Iridium serves as a universal marker for all nuclei in the tissue images, regardless of cell type. Therefore, even if a few cells are not captured by the combined channels used for the cytoplasm and cell membrane, they can still be segmented through the nuclear channel marked by Iridium.

From the Allen Institute’s WTC-11 hiPSC Single-Cell Image Dataset [19], we obtained “full-field” cell culture images containing multiple 3D cells, which had been used to derive all the single-cell images in this dataset. Each image has channels for cell membrane (labeled “CMDRP”) and nuclear (“H3342”) markers, respectively, as well as an additional channel for a tagged marker for one of 25 cellular structures. (They also contain channels labeled “SEG_Memb” and “SEG_DNA” but is it important to note that these are not from human-provided segmentations but rather just contain the output of their segmentation algorithm). We selected five datasets that contain different cytoplasmic structure labels (see Supplementary Table 4 for details). From each of these, we randomly picked five images. The reason for this selection is to obtain a diverse range of cell culture images with different markers, thereby testing and evaluating the generalizability of our pipeline.

### Forming 3D cell candidates

The initial step in our pipeline is carried out (separately) for each of the three axes of the input 3D image. It consists of first applying a given 2D segmentation method to all 2D slices along a given axis, to produce *2D cell regions*. The overlap between 2D cell regions in adjacent slices is then measured using the Jaccard Index (JI). Specifically, the algorithm sequentially considers each 2D cell region in the current slice (referred to below as the query 2D region) and identifies the 2D cell region in the next slice that has the highest JI (if there are any). If that JI is greater than a given threshold, that overlapping region is merged with the query 2D cell region to form a *3D cell candidate* with a unique index (Algorithm 1). A merged overlapping 2D cell region cannot be selected again. This produces 3D cell candidates from each axis, each with a unique index.

It’s important to note that this JI threshold-based matching might lead to elongated stacks with identical index values along a corresponding axis. Even with large overlap, 2D segmented cells in adjacent slices might actually belong to different 3D neighboring cells. This occurs especially frequently in tissue images, where cells are densely packed across all three axes. We address this concern in the subsequent step.

#### Algorithm 1

Forming 3D cell candidates

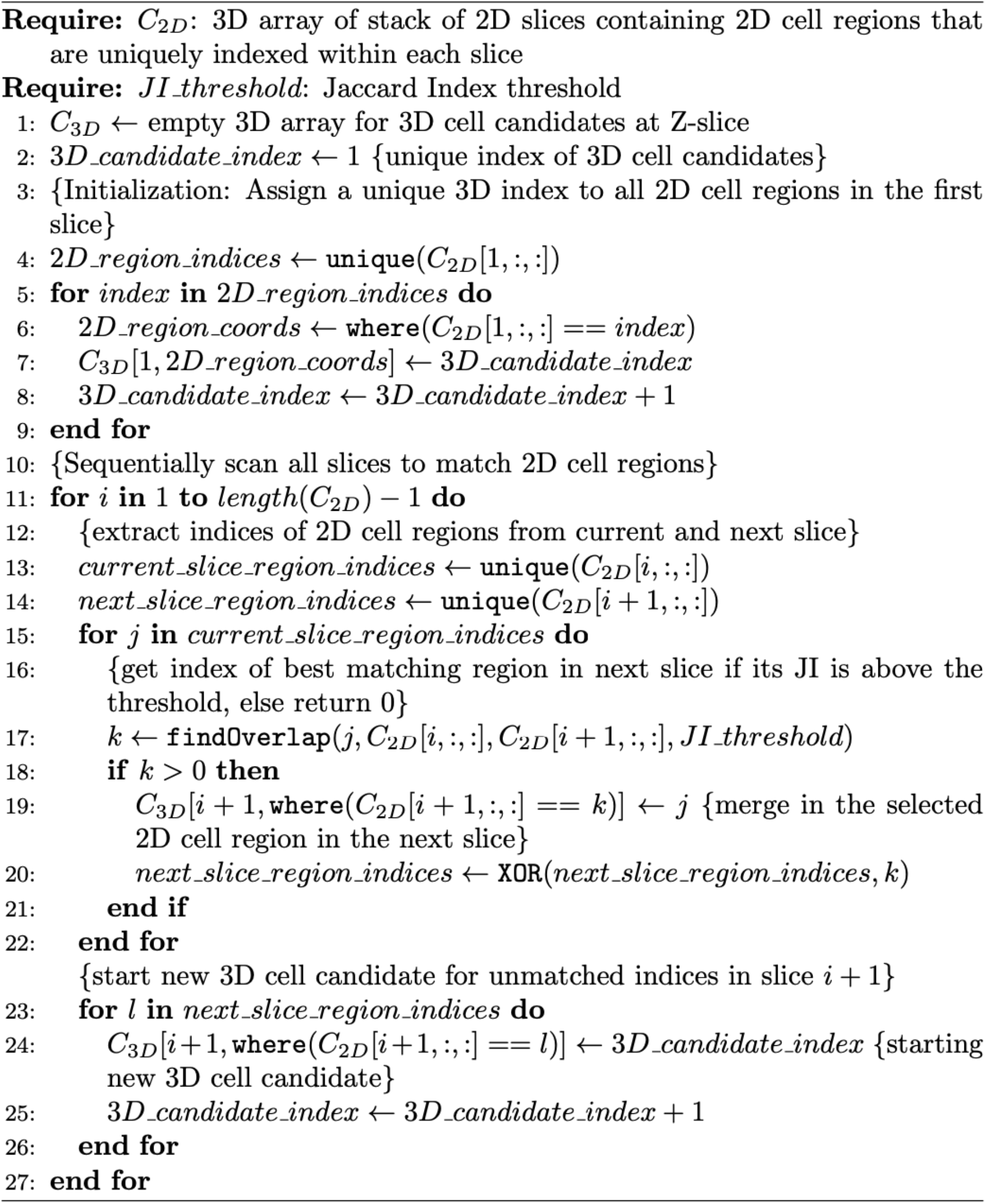

### Matching 3D cell candidates from 3 axes

The next step produces a unified 3D segmentation by integrating the 3D cell candidates from all three axes. We assume that, while creating 3D cell candidates from segmentations along a single axis may be expected to yield errors, combining segmentations from all three axes will help to rectify them.

This approach combines the 3D cell candidates derived from the three axes by assigning voxels that have the same tuple of indices in all three axes to a *3D cell fragment* (Algorithm 2). We also considered that such fragments shorter than three voxels along the z-axis are biologically implausible to correspond to cells, as they cannot contain a nucleus without it protruding outside the cell. Such fragments were therefore eliminated.

#### Algorithm 2

Matching 3D cell candidates from 3 axes

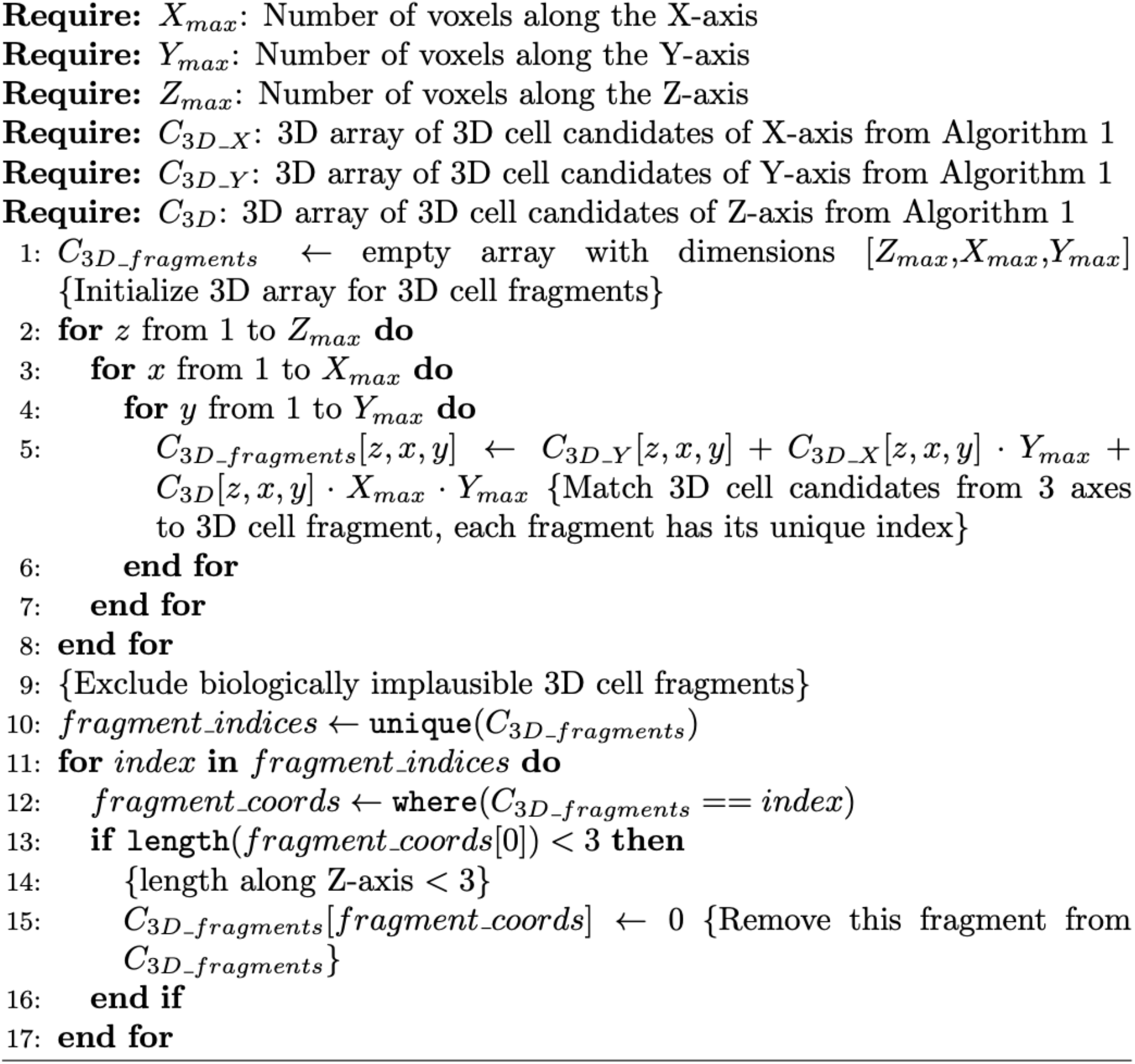

### Refining 3D cell fragments with 3D cell candidates from Z-axis

The 3D cell fragments from the three axes rarely agree perfectly and will almost always be smaller than the actual cell they correspond to. This is because each 3D cell fragment consists only of those voxels that match in all three segmentations, which while they typically agree for large regions of each actual cell, may disagree along edges. We noted that most imaging techniques capture 2D images along the Z axes, in which case the highest spatial resolution is in the X-Y planes. When this is the case, we have a basis for resolving differences between the 3D cell candidates from the three axes by giving preference to the segmentations resulting from Z-axis slicing.

The linear indexing approach allows for straightforward tracing of any voxel in the final 3D segmentation mask back to the index of the 3D cell candidate on each axis to which it belongs. This significantly facilitates the refinement of the final 3D segmentation by using the 2D segmentations from the Z-axis. We initiated the process (Algorithm 3) by selecting matched 3D cells in descending order of volume, as these represent regions where the three axes align best. For each, we identified its corresponding 3D cell candidate from the Z-axis. We replaced each 2D cell boundary on the Z-axis of a given matched cell with the 2D cell boundary from the same Z-slice in the corresponding 3D cell candidate. This produced *final 3D cell regions*.

This refinement process had the effect of using the X and Y axis segmentations primarily to decide where to separate cells along the Z axis. This is illustrated in Supplementary Figure 1, which shows the intersection between the segmentations from the three axes (the 3D cell fragment) and how this determines the Z-axis extent of the final 3D cell region. By employing this approach, we effectively utilized the segmentation capabilities of each axis and exploited the accuracy of segmentation from the axis with the highest resolution. Of course if the image resolution in all directions is the same, this is not needed.

#### Algorithm 3

Refining 3D cells with 3D cell candidates from Z-axis

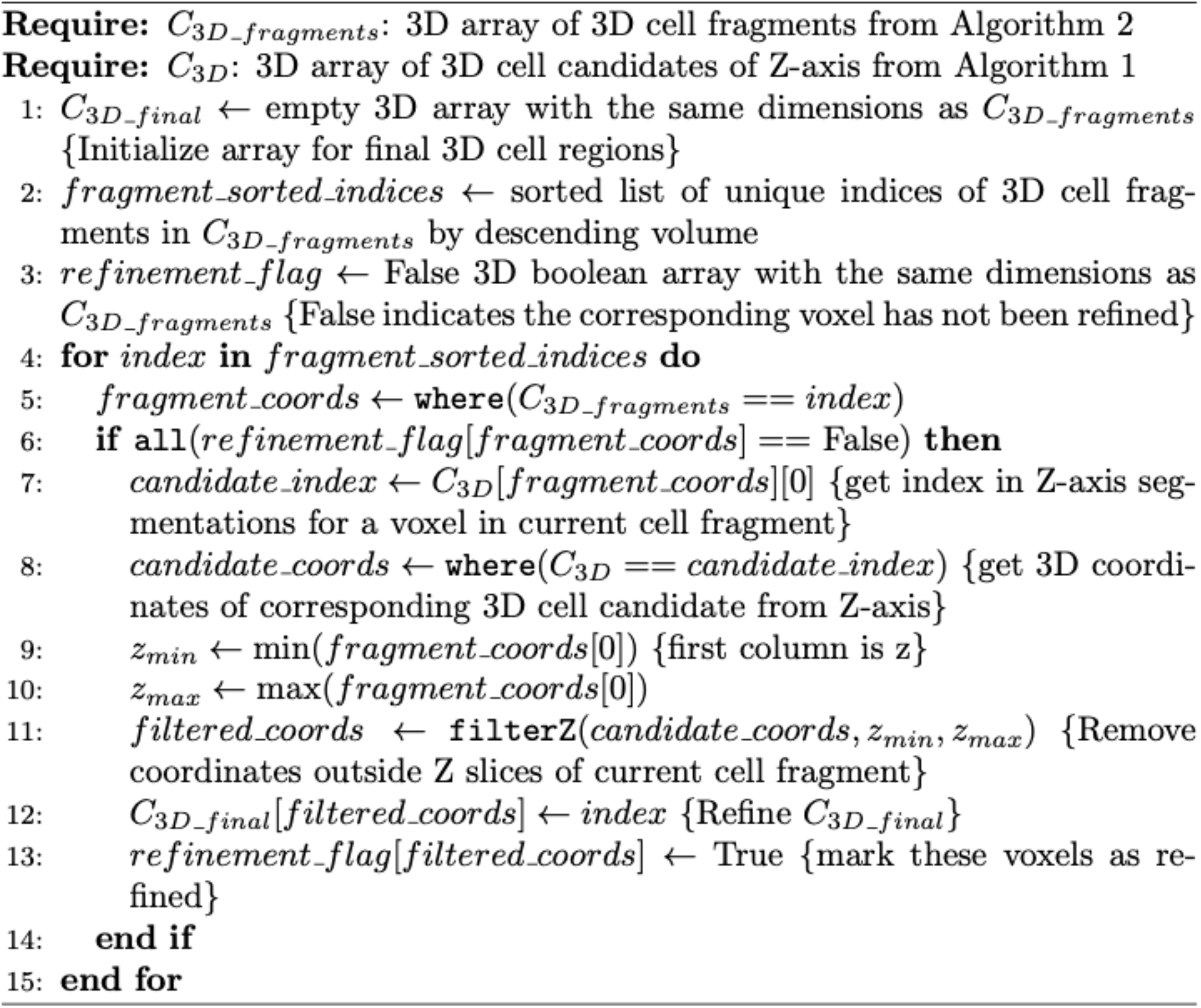

### Inferring 3D segmentation of cell nuclei

For many purposes, 3D cell segmentations are most useful if they include nuclear segmentations. To achieve this, we matched every final 3D cell region from the last step with its corresponding nuclear segmentation mask from the 2D nuclear segmentation along the Z-axis to produce *paired 3D cell and nuclear segmentations* (Algorithm 4). Specifically, we first scanned each 2D slice of every final 3D cell region through the Z-axis and identified its overlapping 2D nuclei from the nuclear mask. During this process, we found that for certain segmentation methods, the nucleus often protruded outside the cell or lay on the cell boundaries. This typically occurred when the methods used separate models for cell and nuclear segmentation. In such case, we trimmed the nucleus to fit within the cell boundaries to maintain the biological plausibility of the cell structure. We also observed that a cell might overlap with multiple nuclei. When this happened, typically there is a dominant nucleus with minimal pixels extending beyond or on the cell boundaries. In contrast, other overlapping nuclei are actually from neighboring cells and thus have a substantial portion outside this cell. Thus, we retained the nucleus with the smallest fraction of protrusion beyond and on the cell boundaries.

For each 3D cell, the 2D nuclei from multiple Z-axis slices within that cell was assigned the same corresponding cell index, forming a *final 3D nuclear region*. When a 3D cell had no nucleus inside, that cell was removed from the final 3D cell segmentation.

If a given cell segmentation method does not also provide nuclear segmentation, we used the 2D nuclear segmentation from DeepCell as a replacement. This avoided penalizing a potentially good cell segmentation method that does not also produce nuclear segmentations.

For simplicity, we refer to the final 3D cell regions as 3D cells in the following.

#### Algorithm 4

Inferring 3D segmentation of cell nuclei

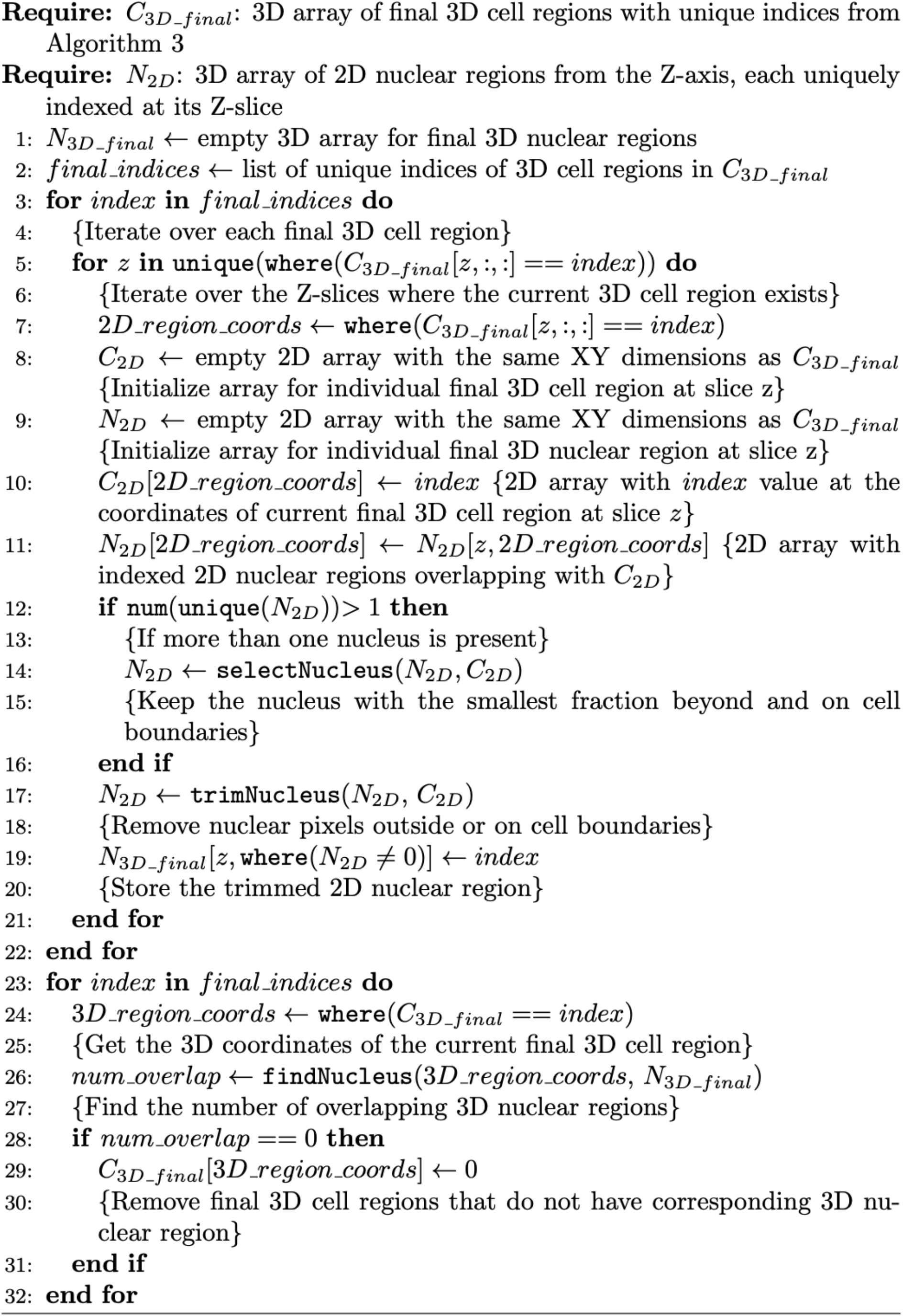

### Visualizing 3D cell segmentation results

We integrated Blender software with 3DCellComposer to visualize 3D cell segmentation masks. For each 3D cell, we used the marching cubes algorithm from the Scikit-learn package of Python to generate a 2D mesh on its surface. This mesh is composed of multiple faces and vertices, which represent the shape and points of the mesh, respectively. We grouped the faces of meshes corresponding to the same 3D cell, allowing Blender to identify them as part of the same structure.

We observed that if the area of a 3D cell in the first z-slice is larger than that in the second z-slice (or if the area in the last z-slice is larger than in the second-to-last z-slice), the marching cube algorithm forms a meshed surface inside the cell rather than outside. This introduces a hole in the 3D cell visualization. To ensure a closed surface for every 3D cell, we added a 2D mesh to the first and last z-slices of any 3D cell exhibiting this issue.

We assigned colors to 3D cells with the requirement that neighboring cells not have the same color. Unlike in 2D, four colors are insufficient for this purpose in 3D. The software therefore iteratively uses increasingly larger sets of colors until the neighboring color requirement is met.

### Evaluating 3D cell segmentation results

Beyond the qualitative assessment through visualization, we aimed to quantitatively evaluate the 3D cell segmentation results. The major challenge in doing this is determining the criteria to be used. Traditionally, this is done by comparing to cell segmentations created by human labeling. However, human labeling exhibits variability across different annotators and even from the same annotator over time [21, 22]. Consequently, this can result in annotations that are inferior to what computational algorithms achieve, as demonstrated in our previous work [27]. Therefore, we have previously proposed an alternative approach: measuring various statistical criteria that are expected of good segmentations. Our CellSegmentationEvaluator [27] tool employs a set of innovatively designed metrics based on a series of assumptions about the desirable characteristics of cell segmentation. The most critical one, which is enabled by multichannel imaging, is that properly segmented cells of the same cell type should have similar compositions over available markers. Critically, this avoids the need for time-consuming and error-prone human labeling.

For 3D images, the drawbacks of human labeling are expected to be even greater than for 2D, due to the greater time required and the expected lower accuracy. We therefore expanded CellSegmentationEvaluator to support evaluation of cell segmentation in 3D (see the Supplementary Information for details of metrics and Supplementary Table 5 for a summary). This tool is especially valuable for 3D datasets from emerging methods like 3D IMC, where human segmentations may not be readily available.

While one can assess various aspects of cell segmentation using the individual metrics, we desired a single overall segmentation quality score. We took an empirical approach to defining this. In addition to having results from multiple segmentation methods, we created perturbed images that would be expected to reduce the quality of segmentation. We then z-scored all metrics from all perturbed and unperturbed IMC images and constructed a Principal Component Analysis (PCA) [32-34] model for evaluating multiplexed tissue images. Similarly, we used the z-scored metrics from all hiPSC images to create a PCA model for cultured cell images (this is to take into account the different expectations about the density and arrangement of the cells in the two cases).

PCA begins by computing the covariance matrix 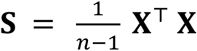, from the z-scored metrics matrix **X**, where *n* is the number of rows in **X**, each row represents a segmentation from one method on one image, and each column a metric. Eigenvalues **λ** and eigenvectors **V** of **S** were then calculated by Singular Value Decomposition (SVD), with the first two eigenvectors **v**_**1**_ and **v**_**2**_ (first two columns of **V**) forming the axes of the first two principal components of the PCA. For each row **x** in **X**, which is a vector of the metrics for a given segmentation method on one image, its first two principal components were computed as PC_1_ = **xv**_**1**_ and PC_**2**_ = **xv**_**2**_.

The final quality score [27] of this mask was then calculated as the weighted sum of PC_1_ and PC_2_:

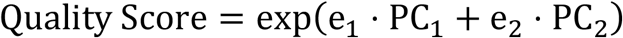

where e_1_ and e_2_ are the proportions of variance explained by first two principal components, respectively, calculated as:

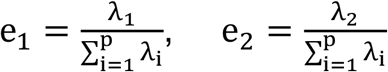

Here, λ_1_ and λ_2_ represent the eigenvalues associated with first two principal components and p is the number of metrics. The denominator sum represents the total variance of **X**. e_1_ and e_2_ emphasize how each component contributes to capturing the variation in the segmentation quality metrics. By applying an exponential function to this summed value to obtain the final quality score, we ensured that the final quality score has a baseline of zero, providing a clear reference for users.

## Results

### 3DCellComposer enables cell segmentation on multiplexed tissue images

We first evaluated the performance of 3DCellComposer on three Imaging Mass Cytometry (IMC) images (Supplementary Table 2) [28, 29] available from the HuBMAP [31] portal (https://portal.hubmapconsortium.org/). These were of spleen, thymus, and lymph node, ensuring coverage across different types of organs to test the generalizability of our pipeline on 3D tissue images. Notably, these images have voxels that are 1 micron on the X and Y axes and 2 microns on the Z axis. This anisotropy, along with the densely packed cells in the 3D IMC tissue images, introduces challenges for accurate 3D cell segmentation.

We implemented 6 2D segmentation methods (including DeepCell using either a cell membrane or cytoplasmic channel as input), and used each of these methods to segment 2D slices from the X, Y, and Z axes of each 3D IMC image (the 2D methods were configured with appropriate free parameters, see Supplementary Table 6). These segmentation outputs were then processed through our 3DCellComposer pipeline to generate 3D cell segmentations. For comparison, we also used the Cellpose 2D to 3D version, and two direct 3D segmentation methods, ACSS and 3DCellSeg.

Example 3D segmentations for a region of one IMC image are shown in Figure 2. This region contains densely packed lymph node cells. We observed that 3DCellComposer with DeepCell segmented the 3D cells in this area with high accuracy and coverage. In contrast, 3DCellComposer with CellProfiler segmented 3D cells with rough boundaries and missed many cells. Meanwhile, Cellpose 2D to 3D segmented only a few 3D cells, and 3DCellSeg tended to excessively merge neighboring cells.

**Figure 2.**
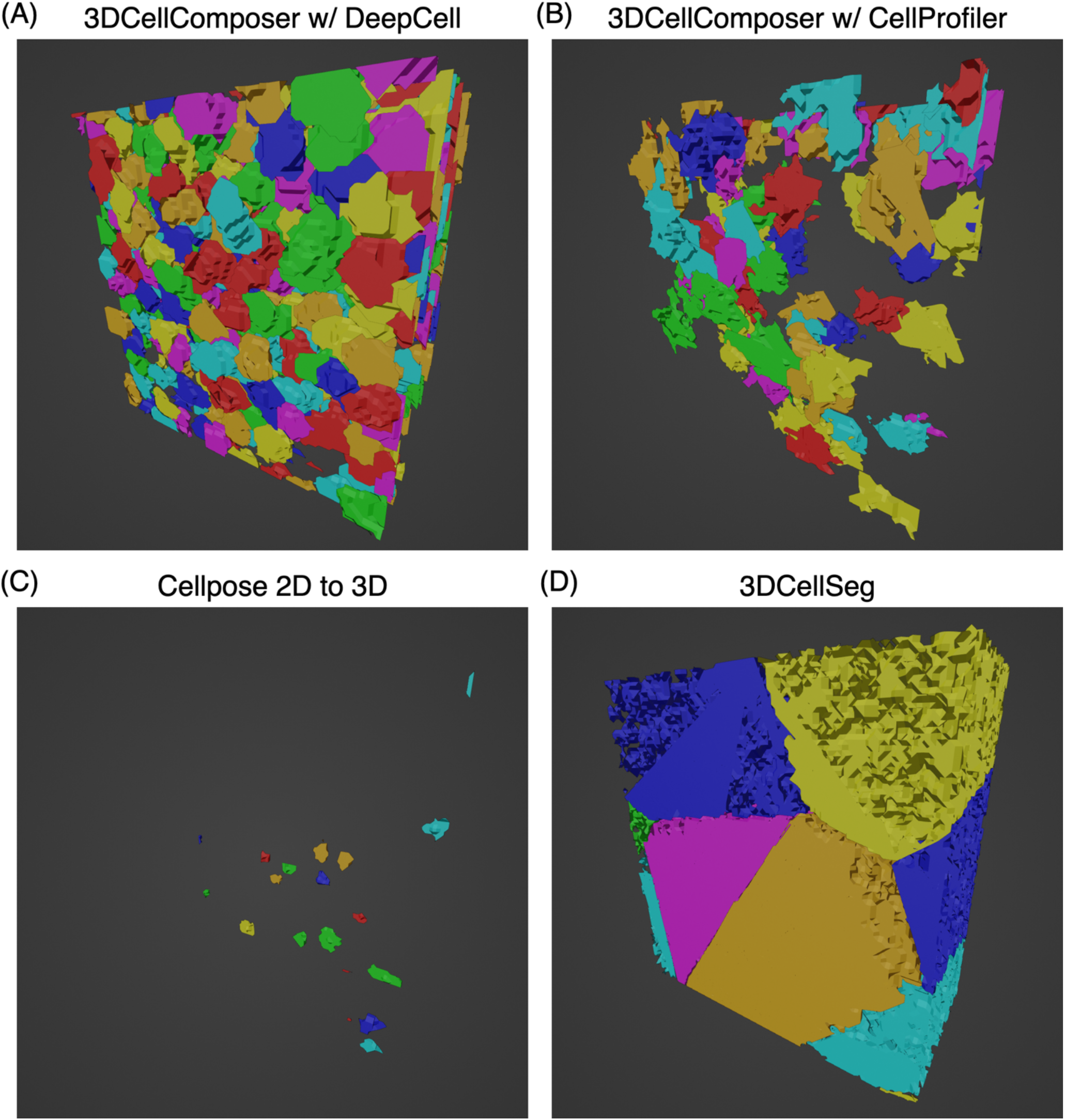
Comparison of 3D cell segmentation using different methods on the Spleen IMC image. Panels A to D display the segmentation outcomes obtained from each respective method. Visualization was carried out using Blender software, utilizing mesh output from 3DCellComposer. To ensure visual distinction, adjacent 3D cells are colored differently.

Examination of the number of cells produced by each method at each step of the process (Supplementary Table 7) reveals that a major reason for the poor performance of all methods except DeepCell is the reduced number of cells detected in 2D slices, especially in the slices along the Z axis. This contributes to the low numbers of 3D cell candidates constructed, since missing cells in one slice inhibits joining of cells in adjacent slices.

While these assessments give a preliminary idea of the characteristics of each method, we proceeded to perform a quantitative evaluation using a 3D extension of our previously described CellSegmentationEvaluator tool [27] (see Methods).

To calibrate our cell segmentation metrics and assess the robustness of segmentation methods against image perturbation, we created three modified image sets from the original IMC images. For these sets, Gaussian noise with a mean of zero was added to each voxel’s intensity. The standard deviation of the noise varied across sets, with values of 5, 10, and 15 voxel intensity, respectively. Notably, a standard deviation of 15 corresponds to approximately 5% of the maximum voxel intensity of the original IMC images. In principle the perturbed images would yield poor segmentations and allow identification of how the metrics responded. To identify key modes of variation among these metrics, we trained a Principal Component Analysis (PCA) model on metrics resulting from all methods applied to all original and perturbed 3D IMC images. While the exact model would depend to some degree on the particular images used for training, the perturbation scheme ensures that the model captures variation across a range of quality. Note that this means that comparison of different methods on the same image should be reliable. Of course, a new PCA model can always be trained on new and perturbed images if desired.

The contributions of each metric to the first two principal components are shown in Supplementary Figure 2. Consistent with our previous study, the first principal component axis reflects the overall segmentation quality, combining all of the metrics except the average cell size but primarily reflecting the measures of marker consistency (which reflect how well the segmented cell boundaries align with the actual cell boundaries). The second principal component indicates the overall coverage of segmentation, primarily reflecting the number of cells and fraction of image covered by cells. A well-performing method should score high on both PC coordinates, as it is expected to perform well across all metrics.

Figure 3A shows the average of PC1 and PC2 for each method on the original IMC images and on the artificially perturbed images. We found that 3DCellComposer, using the DeepCell 2D segmentation model with either cell membrane or cytoplasmic input, achieved the best performance in segmenting 3D cells from IMC tissue images, yielding high values in both quality (PC1) and coverage (PC2), along with high robustness under Gaussian noise perturbation. The performance was marginally better using the cell membrane input. This is consistent with the conclusion from the previous 2D segmentation study, which found that DeepCell performs better using a cell membrane marker than a cytoplasmic marker [27].

**Figure 3.**
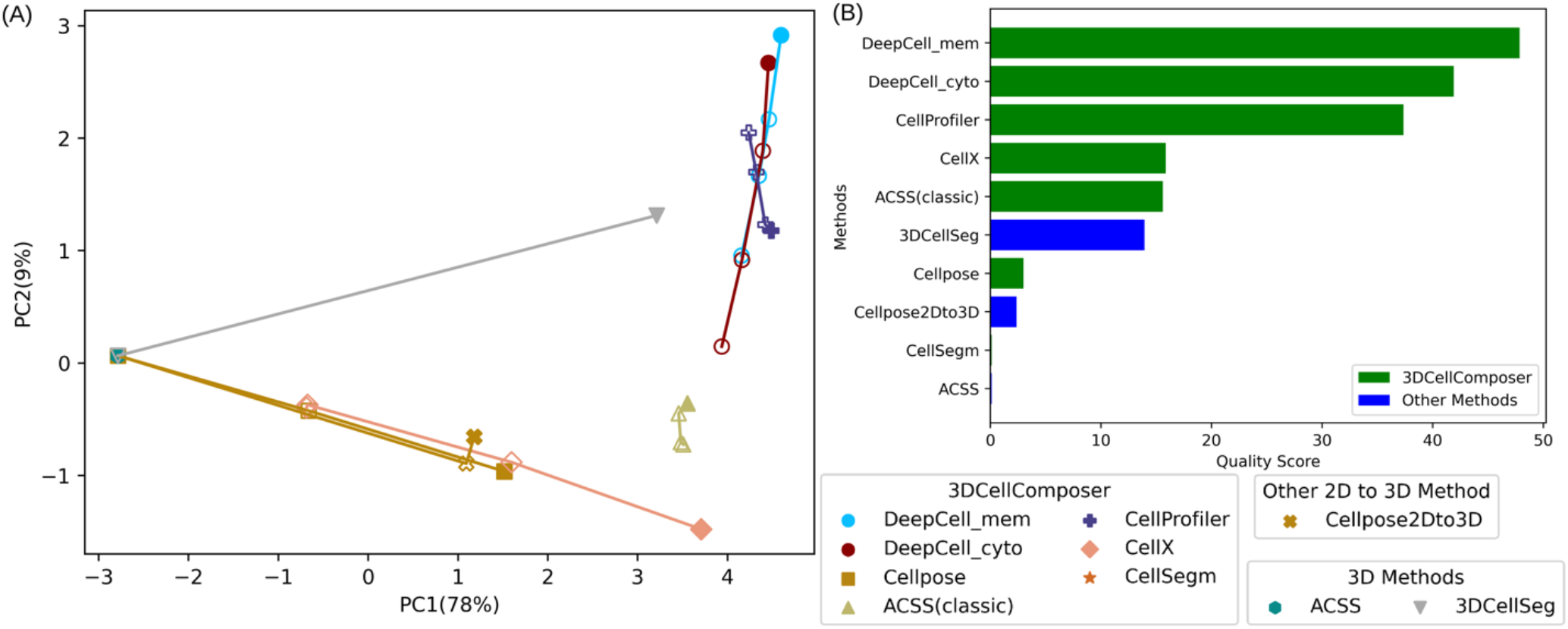
Evaluation of 3D cell segmentation quality on 3D multiplexed tissue images without manual annotations. (A) The top two principal components (PCs) of the segmentation quality metrics for 3DCellComposer, based on different 2D segmentation methods across three organs. The percentages next to PC denote the variance that each corresponding PC accounts for. These results are compared against Cellpose 2D to 3D and two direct 3D methods ACSS, and 3DCellSeg. Each method is represented by a unique color and distinct marker, while filled markers correspond to original images. Hollow marker trajectories illustrate varying levels of Gaussian noise perturbation, arranged in ascending order from low to high. The suffixes “mem” and “cyto” after DeepCell denote the cell membrane and cytoplasmic input channels, respectively. (B) A ranking of 3D segmentation methods by overall segmentation quality score. The green bars are 2D methods through 3DCellComposer whereas blue ones are other methods.

We also found that 3DCellComposer combined with Cellpose did not produce acceptable 3D segmentations, with both PC1 and PC2 being low. This may be due to the fact that, as indicated by our previous study, Cellpose did not perform well on 2D IMC images [27] (this is not surprising in that DeepCell training included IMC images, while Cellpose training did not). It also explains the poor performance of Cellpose 2D to 3D seen in Figure 2.

3DCellComposer displayed varying segmentation capabilities when combined with traditional 2D cell segmentation methods. 3DCellComposer paired with CellProfiler not only demonstrated good segmentation quality (high PC1) and coverage (high PC2) but also performed well under noise perturbation. However, the ACSS classic version and CellX, while acceptable in segmentation quality (high PC1) and robustness, did not achieve good segmentation coverage (low PC2), reflecting that there are many cells they failed to segment. These series of results are also consistent with the performance of these traditional 2D methods on 2D IMC images [27].

As for two direct 3D cell segmentation methods, 3DCellSeg obtained acceptable coverage (high PC2) but low segmentation quality (low PC1). This is consistent with the example in Figure 2 in which the large cells it produces provide high coverage but do not accurately reflect the real cell boundaries. ACSS failed to segment any cells across 3 tissue images. The performance of these two 3D cell segmentation models is consistent with our premise that models trained directly on 3D cell segmentation data might have limited applicability.

To facilitate easy and straightforward comparisons of 3D cell segmentation quality, we calculated the final quality score (see Methods for calculation details) for various methods for 3D IMC images (Figure 3B). The rankings are consistent with the conclusions drawn from Figure 3A.

### 3DCellComposer optimizes segmentation criterion for different 3D images

The most important part of the 3DCellComposer algorithm is determining how to combine the cells found in the 3 different 2D views. This is controlled by a free parameter, a Jaccard Index (JI) threshold, that determines whether two 2D cells in adjacent slices along the X, Y, or Z axis are considered to belong to the same 3D cell. The principle is that if segmented cells in adjacent slices have good overlap, they are likely to be part of the same cell (see Methods for detailed description of this process). 3DCellComposer automatically chooses this threshold for a given method and image to optimize the 3D cell segmentation performance (based on the metrics). Below we describe the process and its effectiveness in detail.

For illustration, we selected a range of Jaccard Index (JI) threshold values (from 0.0 to 0.7 with 0.1 intervals) for whether to match 2D cells in adjacent slices, spanning from very loose to very strict criteria. We applied each threshold with each of the 2D segmentation models to example IMC images (overriding the automatic threshold selection).

We then trained a PCA model using our metrics from all segmentation results from 2D segmentation models across the range of JI settings, along with segmentations from Cellpose 2D to 3D and the direct 3D segmentation methods. This comprehensive model was then used to measure segmentation quality as a function of JI.

Supplementary Figure 3 shows results (using DeepCell with cell membrane input) for three measures for three 3D IMC images: the total number of pairs of matched 2D cells in adjacent slices across all 3 axes, the total number of 3D cells produced, and the segmentation quality score. As noted in the caption, the final number of 3D cells is in rough agreement with expectations given the image dimensions and the typical cell size variation.

We first observed that the quality score increases monotonically as the JI threshold decreases until at least 0.1. In other words, a looser criterion for potentially matching 2D cells between adjacent slices generally leads to higher 3D segmentation quality. This trend may be due to the relatively large voxel size (1×1×2 microns) of these 3D IMC images. Compared to other multiplexed imaging modalities like CODEX [35], which typically have three times higher resolution, this relatively low resolution along the Z-axis leads to more considerable shifts in cell shape between 2D cells belonging to the same 3D cell in adjacent slices (and thus requires a looser criterion to match them). This is supported by the distributions of the number of pairs of matched 2D cells and of matched 3D cells, which both rise with decreasing JI criterion. The increased number of 3D cells results in higher segmentation quality scores.

It is important to note that this optimized low JI threshold setting for DeepCell/3DCellComposer on 3D IMC images need not be universally applicable to other 2D segmentation methods or other imaging modalities.

Interestingly, for two example images the JI threshold yielding the highest quality score was at 0.1. Our choice to incorporate automatic determination of JI threshold for each image and segmentation method accounts for variation from image to image and tissue to tissue.

### 3DCellComposer accurately segments 3D cell culture images

To get an idea of the generalizability of 3DCellComposer, we applied it on 3D images of cell cultures. As described in Methods, we downloaded 25 3D hiPSC cell culture images from the Allen Institute for Cell Science, five images each that are labeled for alpha-actinin, microtubules, Golgi apparatus, lysosomes, and mitochondria. All 25 images include a nucleus marker channel and a cell membrane marker channel.

On this dataset, we tested 3DCellComposer in combination with 2D cell segmentation models DeepCell with cell membrane input and Cellpose (both of them are deep learning models), and compared the results with direct 3D segmentation method ACSS (Figure 4). Since the 3D cell segmentation model in ACSS was trained on a subset of 3D hiPSC cell culture images with a human-in-the-loop approach, we anticipated that ACSS would serve as a strong benchmark for evaluating the performance of 3DCellComposer.

**Figure 4.**
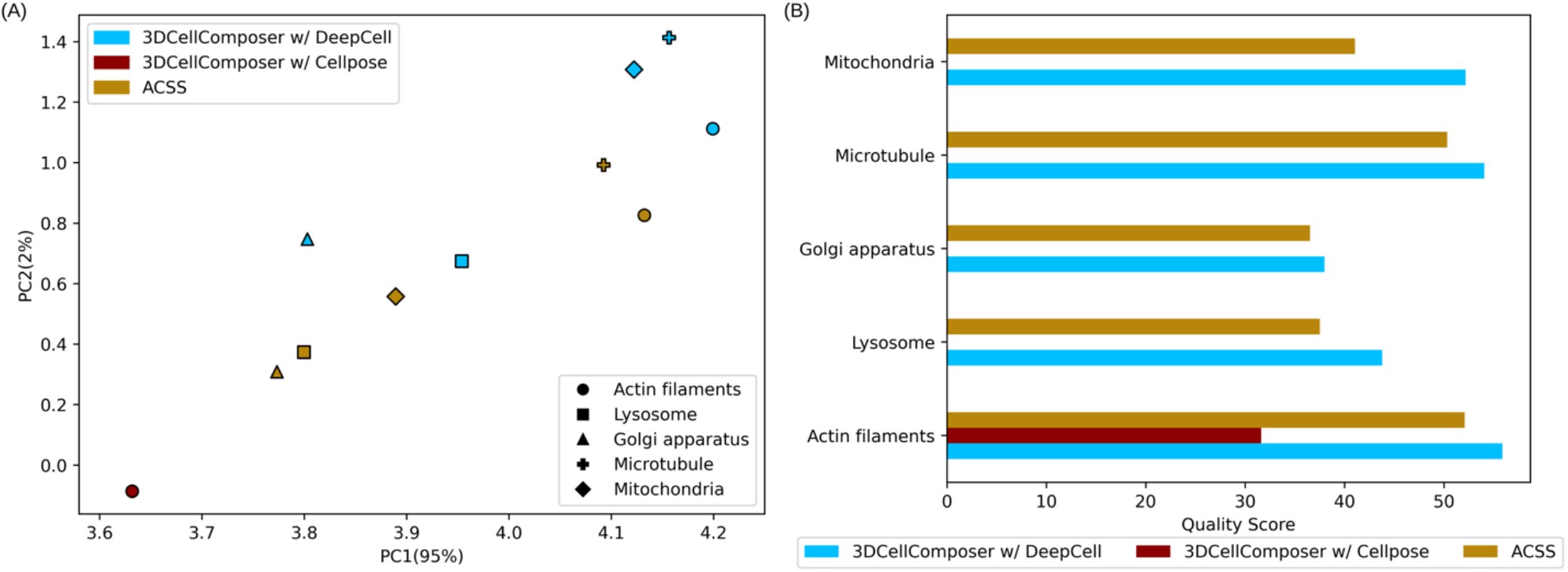
(A) The top two principal components are illustrated for 3DCellComposer and ACSS on 3D hiPSC cell culture images for five cellular markers. Each method is denoted by a unique color, and distinct marker shapes correspond to the five cellular markers. (B) A ranking based on overall quality scores for 3D segmentation methods is presented for the five cellular markers.

For each method, we employed the same segmentation and evaluation protocols as those used for the 3D IMC images, and we visualized the top two principal components (Figure 4A and Supplementary Figure 4) along with the final quality scores (Figure 4B).

We found that 3DCellComposer paired with DeepCell exhibited consistently high segmentation performance across the five image subsets with distinct cytoplasmic markers. ACSS also demonstrated strong performance as expected. For every subset, the combination of 3DCellComposer with DeepCell outperformed ACSS’s 3D segmentation. This not only emphasizes the broad applicability of 3DCellComposer but also the value of our segmentation metrics for evaluating alternative methods even for 3D cell datasets that have a specialized 3D segmentation model like ACSS.

On the other hand, when 3DCellComposer was paired with Cellpose, it only produced reasonable segmentation for the alpha-actinin subset and failed to generate meaningful segmentation for other subsets. This observation underscores the fact that, depending on the input channel for segmentation, a method might work well for one marker but not for another, even if they are from the same cell type and imaging modality. This highlights the need for flexibility in 3D cell segmentation, which 3DCellComposer provides by being able to select from multiple pre-trained 2D segmentation models.

Furthermore, our analysis indicates that 3DCellComposer paired with DeepCell was more resilient than ACSS when processing perturbed images affected by Gaussian noise (Supplementary Figure 5). This suggests that 3DCellComposer can inherit the robustness from 2D cell segmentation models, such as DeepCell in this case, and that 3D deep learning methods such as ACSS that are trained with a specific voxel resolution may be less able to generalize to other resolutions.

Similar to the approach with 3D IMC images, we also examined JI optimization on the hiPSC dataset (Supplementary Figure 6; as noted in the caption, the number of final 3D cells is again in approximate agreement with expectation). In contrast to 3D IMC images, hiPSC cells exhibited significant overlap between 2D cells in adjacent slices. At high JI thresholds, not all adjacent slices were matched, resulting in 3D cells being split and a higher total 3D cell count. As the JI threshold is lowered, the number of matched slices increased, reducing the splitting of 3D cells and resulting in a lower total count of final 3D cells. Very low values may result in overmerging of 3D cells. The quality score values allow choice of the optimal JI, which varied for different images but was typically in the range of 0.4-0.5. This is much higher than optimal for the IHC images, emphasizing the value of our including automatic JI optimization for every 2D segmentation model applied in 3DCellComposer.

## Discussion

As advances in 3D high-content imaging techniques continue to progress, the need for effective 3D cell segmentation has become increasingly important. We have demonstrated here that 3DCellComposer can leverage any well-trained 2D cell segmentation model and perform 3D cell segmentation tasks across two different imaging modalities and voxel resolutions. Our pipeline succeeded in segmenting 3D cells in IMC multiplexed tissue images from three different organs, a task that other 3D cell segmentation methods failed to accomplish. Integration with Blender allows quick visual assessment of the quality of segmentation (Figure 2). We also enhanced our segmentation evaluation tool, CellSegmentationEvaluator, to support 3D segmentations and integrated it into 3DCellComposer. This allows for a direct comparison of 3D segmentation results without the need for manual annotation (Figure 3). Finally, our software outperformed the ACSS model, a direct 3D segmentation model, in segmenting hiPSC cell culture images labeled for various cellular structures, even though the ACSS model was originally trained on images from the same collection (Figure 4). This result further underscores the effectiveness and broad applicability of our software.

Our pipeline is designed to resolve the existing gap between the rise of 3D multiplexed tissue imaging and the current shortage of accurate 3D cell segmentation models. 3D multiplexed images are more challenging to prepare and costlier to produce [6, 36]. This limits the number of 3D images produced for a given 3D modality. However, accurate cell segmentation methods, especially those based on deep learning, require a substantial amount of training data. We bridge this gap between availability and necessity by leveraging the generalizability of 2D cell segmentation models and making them suitable for 3D cell segmentation images. Importantly, as 3D multiplexed images often evolve from established 2D imaging modalities (e.g., 2D IMC vs. 3D IMC), a 2D cell segmentation model trained on a particular 2D imaging modality becomes a natural fit for its 3D counterpart. This applicability persists even when a 3D modality is in its developmental phase. As a result, our pipeline represents a potentially solution for imaging specialists. Not only does it remove the concerns of cell segmentation during 3D imaging technique development, but it also serves as a strategic guide, directing specialists to prioritize 3D imaging modalities that can benefit from existing, trained 2D models.

Our pipeline provides users with a clear pathway to select the best 3D cell segmentation method for their specific 3D tissue or cell images. From a quantitative perspective, the enhanced CellSegmentationEvaluator evaluates 3D cell segmentation performance without relying on human-provided ground truth. This approach liberates users from burdensome, time-consuming and potentially error-prone manual annotations for newly acquired tissue and cell images.

## Declarations

## Ethics approval and consent to participate

Not applicable

## Consent for publication

Not applicable

## Data availability

- 3DCellComposer software is available at https://github.com/murphygroup/3DCellComposer. It is also available as PyPI package “ThreeDCellComposer” (e.g., via *pip*).
- CellSegmentationEvaluator is available at https://github.com/murphygroup/CellSegmentationEvaluator. It is also available as PyPI package “CellSegmentationEvaluator” (e.g., via *pip*).
- All data used for this work are available as a Reproducible Research Archive, https://github.com/murphygroup/ChenMurphy3DCellComposerRRA.

## Author contributions

H.C. conceived the study, performed computational analysis, and wrote the manuscript. R.F.M. conceived the study, supervised the study, and wrote the manuscript. All authors approved the content of the manuscript.

## Acknowledgements

We thank Matthew Ruffalo for helpful discussions.

## Authors’ information (optional)

Not applicable

