## Supplementary Information for "3DCellComposer - A Versatile Pipeline Utilizing 2D Cell Segmentation Methods for 3D Cell Segmentation"

### Description of individual cell segmentation metrics

We defined 14 metrics to evaluate the performance of a single 3D segmentation method without requiring a reference segmentation. These are of two types: metrics that assess the coverage of a segmentation mask on the image, and metrics that measure various types of uniformity at the voxel and cell levels on multiplexed images. Each metric is derived under an assumption based on general concepts from cell biology. They are described in the following and summarized in Supplementary Table 6.

*Coverage Metrics*

The general assumption of coverage metrics is that a tissue image should be mostly occupied by cells except for readily detectable volumes corresponding to ducts, blood vessels, or tissue borders. Therefore, we assume that a good segmentation should generate a mask that covers most volumes of the tissue in an image.

**Number of cells per 100 cubic microns (NC)**

The principle is that cell density is a measure of the quality of both the image and the segmentation method. The voxel size in cubic microns varies in different image modalities. To calculate the density, voxel sizes in the X, Y and Z dimensions are directly obtained from OME-TIFF metadata. The metric is defined as

$$\text{NC} = \frac{n_c}{s_p \times n_p} \times 100$$

where $n_c$ is the number of cells in the cell mask, $s_p$ is the size of one voxel in cubic microns, $n_p$ is the total number of voxels in the corresponding image.

To further assess the coverage of the segmentation mask, each image was divided into two distinct regions: the foreground and the background. The foreground includes the tissue volumes, which are of primary interest, while the background consists of all volumes outside the tissue boundary. To achieve this, we adopted the 2D foreground-background separation method developed for 2D multiplexed tissue images in our previous study [1]. We applied this method to each Z-slice of the 3D multiplexed image, stacking the results to create the final binary foreground-background separated 3D image.

Given the binary foreground-background separation, we designed the following three coverage metrics with the general assumption that an accurate segmentation mask will cover most tissue volumes (i.e., the foreground) but few background volumes.

**Fraction of image foreground occupied by cells (FFC)**

$$\text{FCF} = \frac{a_{cf}}{a_f}$$

where $a_{cf}$ is the volume of a cell mask in the foreground and $a_f$ is the volume of the foreground.

**Fraction of image background occupied by cells (FBC)**

$$\text{FBC} = \frac{a_{cb}}{a_b}$$

where $a_{cb}$ is the volume of a cell mask in the background and $a_f$ is the volume of the background.
1 minus FBC is used as the metric, so that larger values indicate better segmentation.

**Fraction of cell mask in foreground (FCF)**

$$\mathrm{FCF} = \frac{a_{cf}}{a_c}$$

where $a_{cf}$ is the volume of a cell mask in the foreground and $a_c$ is the volume of the cell mask.

*Homogeneity Metrics*

To measure the homogeneities of voxel intensities and integrated 3D cell intensities from a segmentation method, we developed another category called homogeneity metrics.

At the voxel level, the principle is that the voxels outside of the cells but still within the image foreground should be similar in protein composition (i.e., should consist of extracellular matrix of similar composition). The coefficient of variation (CV) of the foreground voxels outside the cells for each channel was calculated and then the average CV across all channels was taken.

**Average CV of foreground voxels outside the cells (ACVF).**

$$\mathrm{ACVF} = \frac{1}{n_h}\sum_{i=1}^{n_h}\frac{\sigma_{if}}{\mu_i}$$

where $n_h$ is the total number of image channels, $\sigma_{if}$ is the standard deviation of voxel intensities of the $i^{th}$ channel in the foreground outside the cells, $\mu_i$ is the mean of all voxel intensities in the $i^{th}$channel.
The reciprocal of ACVF+1 is used as the metric, so that higher values indicate better segmentation and its upper bound is 1.

We applied the Principal Component Analysis (PCA) on the matrix of all foreground outside-the-cell voxel intensities across all channels after z-score standardization on each channel was applied and the fraction of variance explained by the first principal component was calculated. A higher value of the fraction stands for a more conserved relationship across channels in the foreground outside the cell volumes.

**Fraction of first PC of foreground voxels outside the cells (FPCF).**

$$\text{FPCF} = \frac{\lambda_{f1}}{tr(\Sigma_f)}$$

where $\lambda_{f1}$ is the variance explained by the first principal component (PC) across channels for the foreground voxels outside the cells, $\Sigma_f$ is the covariance matrices from PCA analysis on the foreground voxels outside the cells across all channels, $tr(\square)$ is trace calculation on a matrix.

At the cell level, the principles are that (a) cell types should be roughly similar in composition across all channels, (b) cell types can be approximated by cell clusters, (c) the average CV across all clusters for a given number of clusters is a proxy for similarity in composition within cell types, and (d) averaging the average CV across all clusters across different numbers of the cluster is also a proxy for similarity in composition within cell types. Cell types were defined using KMeans clustering performed on the mean cell intensities across all channels after applying z-score standardization for each channel. Since the number of cell types in an arbitrary image is unknown, K, the number of clusters, was varied from 1 to 10. Note that the KMeans clustering was applied on the cell mask to define the cell type, which generated cell type labels of each number of clusters used in the following metric calculation for each cellular component mask.

For the clusters from each K value, the average CV for each cluster across channels was calculated followed by an average weighted by the cluster size. The final metric was calculated as the average of the weighted average coefficient of variation across all clusters over 1 to 10 clusters.

**Average of weighted average CV of cell type intensities over 1-10 clusters (ACVC)**

$$\text{ACVC} = \frac{1}{K_{max} - K_{min} + 1} \sum_{K=K_{min}}^{K_{max}} \left( \frac{1}{n_c} \sum_{k=1}^{K} (CV)_k \cdot n_k \right)$$

$$(CV)_k = \frac{1}{n_h} \sum_{i=1}^{n_h} \frac{\sigma_{ki}}{\mu_{ki}}$$

where $K_{min}$ is the smallest number of clusters, $K_{max}$ is the largest number of clusters, $K$ is the current number of clusters, $n_c$is the total number of cells, $CV_k$ is the coefficient of variation of mean cell intensities in $k^{th}$ cluster, $\sigma_{ki}$ and $\mu_{ki}$ is the standard deviation and the mean of mean cell intensities of $k^{th}$ cluster in $i^{th}$ channel respectively, $n_k$ is the number of cells in $k^{th}$ cluster, $n_h$ is the total number of image channels.

The reciprocal of ACVC+1 is used as the metric, so that higher values indicate better segmentation and its upper bound is 1.

A similar measure was derived using principal component analysis. The first two principles are the same as above but (c) the fraction of variance accounted for by the first principal component of the cells in each cluster is a proxy for similarity in composition of each cell type, and (d) averaging this fraction over different numbers of clusters is also a proxy for similarity of each cell type. We, therefore, applied PCA on the mean cell intensities across z-score standardized channels of each number of cluster and calculated the average fraction of variance accounted for by the first principal component across all numbers of clusters. The output vector was averaged over different K values to get the final metric.

**Average of weighted average fraction of the first PC of cell type intensities over 1-10 clusters**

$$\text{FPCC} = \frac{1}{K_{max} - K_{min} + 1} \sum_{K=K_{min}}^{K_{max}} \left( \frac{1}{n_c} \sum_{k=1}^{K} \frac{\lambda_{k1}}{tr(\Sigma_k)} \cdot n_k \right)$$

where $K_{min}$ is the smallest number of clusters, $K_{max}$ is the largest number of clusters, $K$ is the current number of clusters of choice, $n_c$ is the total number of cells, $\lambda_{k1}$ is the variance explained by the first principle component in $k^{th}$ cluster, $\Sigma_k$ the covariance matrices from PCA on mean cell intensities across all channels in $k^{th}$ cluster, $n_k$ is the number of cells in the current cluster.

We also incorporated the Silhouette score as a metric to indicate the similarity of composition across cell types. Since the Silhouette score equals 1 when there is only one cluster, we started with two clusters to calculate this metric.

**Average of Silhouette score of clustering over 1-10 clusters (AS)**

$$\text{AS} = \frac{1}{K_{max} - K_{min} + 1} \sum_{K=K_{min}}^{K_{max}} \left( \frac{1}{n_c} \sum_{i=1}^{n_c} \frac{b(i) - a(i)}{\max\left(a(i), b(i)\right)} \right)$$

$$\mathrm{a}(i) = \frac{1}{|K_i| - 1} \sum_{j \in K_i, i \neq j} d(i,j)$$

$$\mathrm{b}(i) = \min_{l \neq i} \frac{1}{|K_l|} \sum_{j \in K_l} d(i,j)$$

where $K_{min}$ is the smallest number of clusters, $K_{max}$ is the largest number of clusters, $K$ is the current number of clusters of choice, $a(i)$ is the average distance between cell $i$ and all the other cells $j$ in the cluster $K_i$ to which cell $i$ belongs, $b(i)$ is the minimum average distance from cell $i$ to all cells $j$ in all clusters $K_l$ to which $i$ does not belong, $n_c$ is the total number of cells.

Besides the homogeneity at the voxel and cell levels in composition across image channels, we also assume that properly segmented images should contain cells of similar sizes.

**Coefficient of variation of cell size (CSCV)**

$$\text{CSCV} = \frac{\sigma_s}{\mu_s}$$

where $\mu_s$ represents the average size in voxel across all 3D cells, and $\sigma_s$ stands for the standard deviation of all cell sizes.

The reciprocal of CSCV+1 is used as the metric, so that higher values indicate better segmentation and its upper bound is 1.

We also developed a metric that penalizes over-segmentation, a more common issue in 3D cell segmentation compared to 2D. This metric regards segmentations that segment larger cells.

**Average cell size weighted by cell size in cubic microns (WACS)**

$$\mathrm{WACS} = \frac{\sum_{i=1}^{n_c}(s_i \cdot s_i)}{\sum_{i=1}^{n_c} s_i}$$

where $s_i$ is the size of $i^{th}$ cell, $n_c$ is the total number of cells. This results in a weighted average cell size metric that gives greater weight to larger cells.

Supplementary Table 1. Summary of segmentation methods

| Method | Dimensions | Category | Input Channels | | | Output | Reference |
|---|---|---|---|---|---|---|---|
| | | | Cytoplasm | Cell Membrane | Nucleus | Nuclear mask* | |
| DeepCell v0.12.6 | 2D | Deep learning | X (DeepCell_cyto) | X (DeepCell_mem) | X | Yes | [2] |
| Cellpose v2.2.2 | 2D | Deep learning | X | | X | Yes | [3, 4] |
| CellSegm | 2D | Conventional | | X | X | No | [5] |
| CellX | 2D | Conventional | | X | | No | [6] |
| CellProfiler | 2D | Conventional | | X | X | Yes | [7, 8] |
| ACSS (classic) | 2D | Conventional | | X | X | No | [9] |
| Cellpose2D-3D | 2D to 3D | Deep learning | X | | X | Yes | [3] |
| ACSS(ML) | 3D | Deep learning | | X | X | Yes | [9] |
| 3DCellSeg | 3D | Deep learning | | X | X | Yes | [10] |

*All methods produce cell masks; this column specifies whether a nuclear mask is also generated.

Supplementary Table 2. Details of 3D IMC datasets. A script to download the images can be found in the Reproducible Research Archive at https://github.com/murphygroup/ChenMurphy3DCellComposerRRA

| Dataset | Tissue | Image Dimensions (ZXY), in voxels of 2 um x 1 um x 1 um |
|---|---|---|
| HBM387.XZWR.467 https://portal.hubmapconsortium.org/browse/dataset/a296c763352828159f3adfa495becf3e | Lymph Node | 48x760x740 |
| HBM778.VHHR.349 https://portal.hubmapconsortium.org/browse/dataset/cd880c54e0095bad5200397588eccf81 | Thymus | 22x597x571 |
| HBM459.CGSD.533 https://portal.hubmapconsortium.org/browse/dataset/d3130f4a89946cc6b300b115a3120b7a | Spleen | 50x514x452 |

Supplementary Table 3. Summary of input channels of IMC for 3D cell segmentation

| Segmentation Input | Marker (Isotope) | Target | Cell type | Source |
|---|---|---|---|---|
| Nucleus | Iridium (Ir191) | nucleus | All cell types | [11] |
| Cytoplasm | Smooth Muscle Actin (SMA) (In115) | smooth muscle actin | smooth muscle cells and myofibroblasts | https://www.uniprot.org/uniprotkb/P63267/entry#subcellular_location |
| | Granzyme B (Tb159) | cytoplasmic granules | cytotoxic T lymphocytes and natural killer (NK) cells | [12] |
| | Myeloperoxidase MPO (Y89) | lysosome | neutrophils and monocytes | https://www.uniprot.org/uniprotkb/P05164/entry#subcellular_location |
| Cell membrane | E-Cadherin / P-Cadherin(La139) | cadherins on cell membrane | epithelial cells | https://www.uniprot.org/uniprotkb/P12830/entry#subcellular_location https://www.uniprot.org/uniprotkb/P22223/entry#subcellular_location |
| | pan Cytokeratin(Pr141) | cadherins on cell membrane | epithelial cells | https://www.uniprot.org/uniprotkb/Q16195/entry#subcellular_location |
| | CD31(Eu151) | cell membrane | endothelial cells and some leukocytes | https://www.uniprot.org/uniprotkb/P16284/entry#subcellular_location |
| | CD45RA(Gd160) | cell membrane | Isoforms of CD45, which is a pan-leukocyte marker | https://www.uniprot.org/uniprotkb/P08575/entry#subcellular_location |
| | CD45RO(Dy162) | cell membrane | Isoforms of CD45, which is a pan-leukocyte marker | https://www.uniprot.org/uniprotkb/P08575/entry#subcellular_location |

Supplementary Table 4. Details of 3D Allen Institute's WTC-11 hiPSC Single-Cell Image Dataset. The images can be viewed at https://www.allencell.org/3d-cell-viewer.html by selecting the protein tag and “Full field” for the image type. A script to download the images can be found in the Reproducible Research Archive at https://github.com/murphygroup/ChenMurphy3DCellComposerRRA

| Datasets (w/ different fluorescence-labeled organelles) | Image Dimensions (ZXY), in voxels of 0.29 um x 0.108 um x 0.108 um | Nuclear channel | Cell membrane channel |
|---|---|---|---|
| Alpha-actinin | 65~70x924x624 | H3342 (Hoechst 33342) | CMDRP (CellMask Deep Red plasma membrane stain) |
| Microtubules | 59~79x924x624 | | |
| Lysosomes | 65~70x924x624 | | |
| Golgi apparatus | 65~75x924x624 | | |
| Mitochondria | 65~70x924x624 | | |

Supplementary Table 5. Summary of the 3D evaluation metrics

| Metric | Metric Abbreviation | Mask(s) to calculate the metrics on |
|---|---|---|
| Number of cells per 100 cubic microns | NC | Cell mask |
| Fraction of image foreground occupied by cells | FFC | Cell mask |
| Fraction of image background occupied by cells | 1-FBC | Cell mask |
| Fraction of cell mask in foreground | FCF | Cell mask |
| Coefficient of variation (CV) of cell size | 1/(CSCV+1) | Cell mask |
| Average cell size weighted by cell size in cubic microns | WACS | Cell Mask |
| Average coefficient of variation (CV) of foreground voxels outside the cells | 1/(ACVF+1) | Cell mask |
| Fraction of first principal component (PC) of foreground voxels outside the cells | FPCF | Cell mask |
| Average of weighted average coefficient of variation (CV) of cluster intensities over number of clusters from 1 to 10 | 1/(ACVC+1) | Nuclear mask (NUC) and cell excluding nucleus mask (CEN) |
| Average of weighted average fraction of the first principal component (PC) of cluster intensities over number of clusters from 1 to 10 | FPCC | Nuclear mask (NUC) and cell excluding nucleus mask (CEN) |
| Average of Silhouette score of clustering over number of clusters from 1 to 10 | AS | Nuclear mask (NUC) and cell excluding nucleus mask (CEN) |

Supplementary Table 6. Parameters for 2D cell segmentation methods

| Method | Free Parameter | Value | |
|---|---|---|---|
| | | For 3D IMC | For 3D hiPSC |
| DeepCell v0.12.6 | Voxel size in micron | 1 (Z-slices)<br>2 (X-slices)<br>2 (Y-slices) | 0.1083 (Z-slices)<br>0.29 (X-slices)<br>0.29 (Y-slices) |
| Cellpose v2.2.2 | model type | cyto | cyto2 |
| | Estimated cell diameter in voxel | 10 (Z-slices)**<br>5 (X-slices)<br>5 (Y-slices) | 100 (Z-slices)<br>37 (X-slices)<br>37 (Y-slices) |
| | Estimated nuclear diameter in voxel | 10 (Z-slices)<br>5 (X-slices)<br>5 (Y-slices) | 100 (Z-slices)<br>37 (X-slices)<br>37 (Y-slices) |
| CellSegm | Minimum and maximum cell diameter | 10 to 40 microns*** | N/A* |
| CellX | Minimum and maximum seed radius | 5 to 10 microns | N/A |
| | Maximum cell length | 50 microns | |
| CellProfiler | Minimum and maximum nucleus diameter | 5 to 10 microns | N/A |
| | Maximum expansion of cell boundary from nucleus | 20 voxels | N/A |
| ACSS (classic) | Minimum cell area | 5 micron$^2$ | N/A |
| Cellpose2D-3D | Anisotropy | 2 | N/A |

* N/A indicates the method was not tested on the dataset
** diameter larger than 10 yielded significantly fewer cells
*** all free parameters in microns were converted when necessary to voxel sizes

Supplementary Table 7. Comparison of cells identified in different stages of 3DCellComposer using various 2D segmentation methods

| Number of cells / Method | DeepCell (mem) | DeepCell (cyto) | Cellpose | ACSS (classic) | CellProfiler | CellX | CellSegm |
|---|---|---|---|---|---|---|---|
| *Lymph Node IMC image* | | | | | | | |
| 2D cells along all slices of Z axis | 264,535 | 256,672 | 43,935 | 88,950 | 128,172 | 62,784 | 501 |
| 2D cells along all slices of Y axis | 1,100,181 | 1,114,245 | 6,410 | 180,177 | 746,580 | 48,011 | 1086 |
| 2D cells along all slices of X axis | 994,395 | 1,019,802 | 9,955 | 175,440 | 731,356 | 51,048 | 974 |
| 3D cell candidates before 3D nuclear matching | 29,303 | 28,204 | 1,644 | 133 | 7,395 | 63 | 11 |
| 3D cell candidates missing nuclei removed | 3 | 13 | 10 | 1 | 1 | 15 | 4 |
| final 3D cells | 29,300 | 28,191 | 1,634 | 132 | 7,394 | 48 | 7 |
| *Thymus IMC 3D image* | | | | | | | |
| 2D cells along all slices of Z axis | 16,861 | 16,761 | 6,452 | 13,812 | 9,141 | 10,690 | 51 |
| 2D cells along all slices of Y axis | 88,816 | 89,463 | 1,409 | 39,345 | 83,188 | 7,987 | 35 |
| 2D cells along all slices of X axis | 81,103 | 80,342 | 2,188 | 43,811 | 63,214 | 8,906 | 19 |
| 3D cell candidates before 3D nuclear matching | 1,383 | 1,331 | 42 | 540 | 596 | 73 | 1 |
| 3D cell candidates missing nuclei removed | 8 | 3 | 1 | 21 | 1 | 29 | 1 |
| final 3D cells | 1,375 | 1,328 | 41 | 519 | 595 | 44 | 0 |
| *Spleen IMC 3D image* | | | | | | | |
| 2D cells along all slices of Z axis | 171,387 | 163,165 | 516 | 42,288 | 83,446 | 20,998 | 221 |
| 2D cells along all slices of Y axis | 59,3312 | 618,965 | 75 | 73,396 | 397,834 | 15,855 | 188 |
| 2D cells along all slices of X axis | 586,648 | 617,517 | 241 | 68,530 | 404,410 | 15,712 | 148 |
| 3D cell candidates before 3D nuclear matching | 21,350 | 18,037 | 0 | 2,065 | 4,440 | 1,048 | 3 |
| 3D cell candidates missing nuclei removed | 14 | 17 | 0 | 34 | 1 | 92 | 2 |
| final 3D cells | 21,336 | 18,020 | 0 | 2,031 | 4,439 | 956 | 1 |

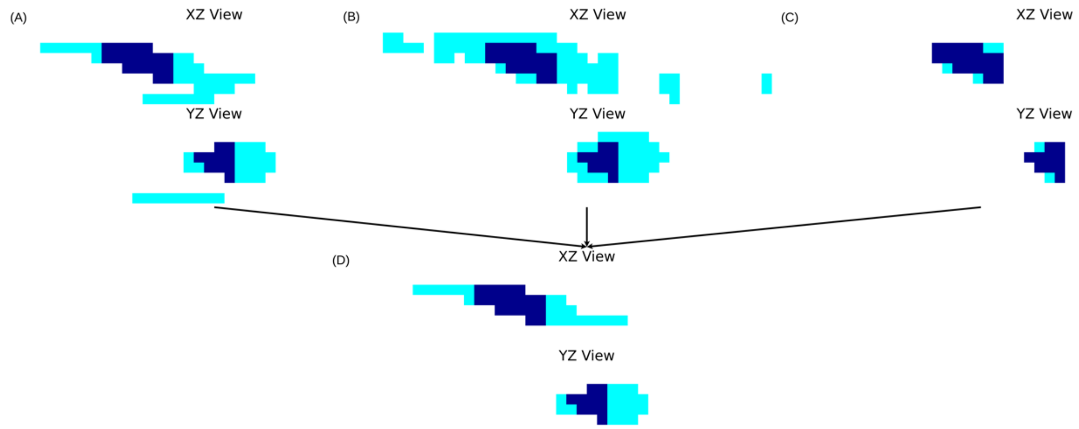


**Supplementary Figure 1** Illustration of the refinement step in 3DCellComposer. X-Z and Y-Z slices containing the centroid of a cell are shown for the 3D cell candidates from slicing along the Z-axis (A), Y-axis (B), and X-axis (C). The common part of different views is colored in dark blue. The linear indexing approach is used to keep the 2D cells from the Z-axis slicing only for those slices in the intersection of all three 3D cell candidates.

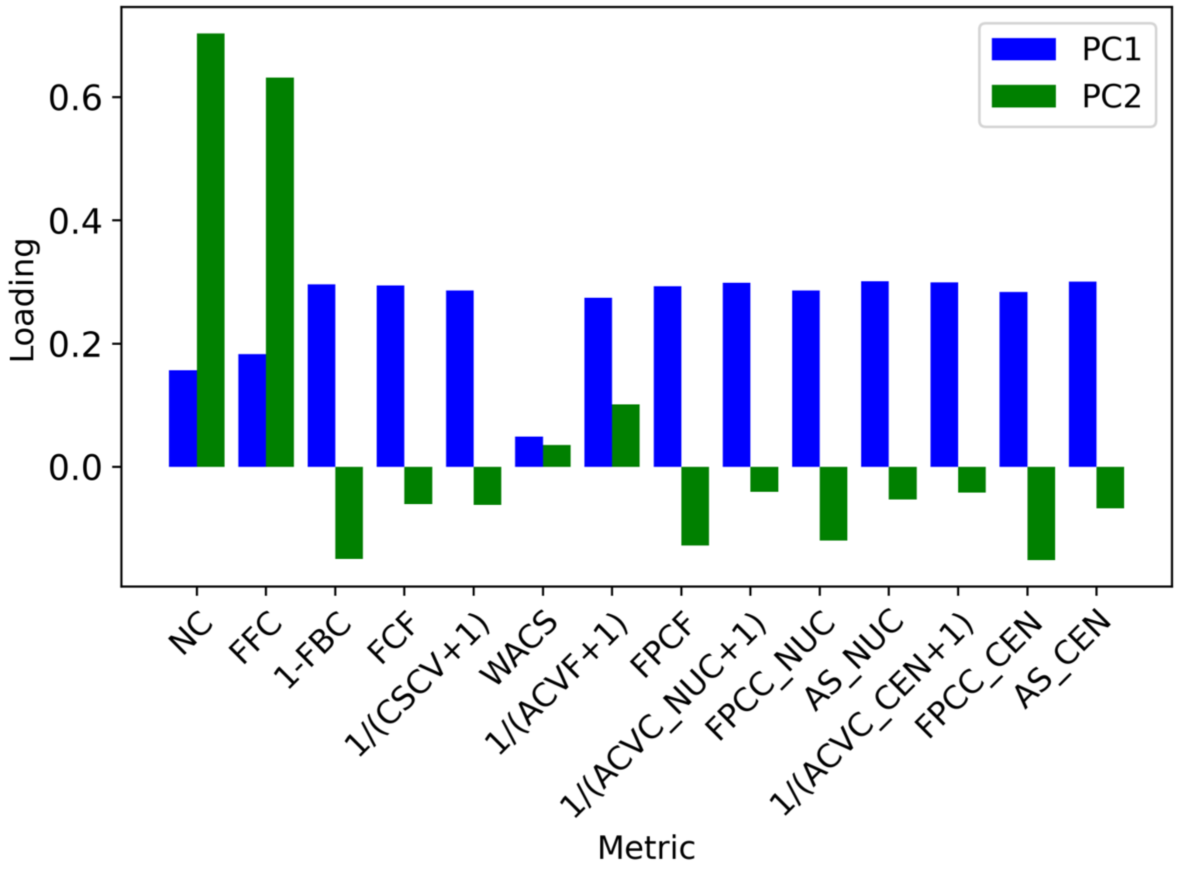


**Supplementary Figure 2** Factor loadings of PCA model trained by all 3D segmentation masks from all methods on 3D IMC images.

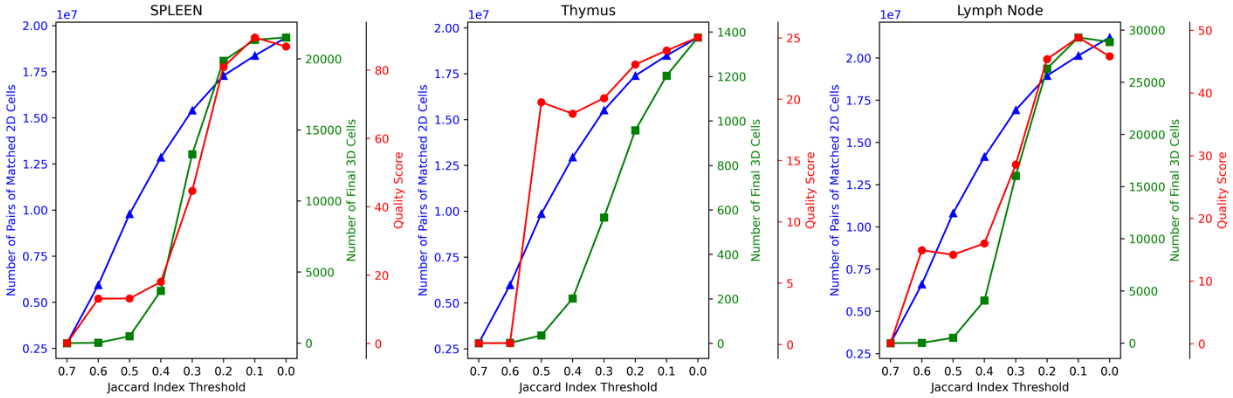


**Supplementary Figure 3** Analysis of the Jaccard Index optimization for 3DCellComposer using DeepCell with cell membrane input on 3D IMC images. The figure compares the total number of matched 2D cell pairs in adjacent slices across all three axes, the total count of segmented 3D cells, and the average quality of overall segmentation. Calculating the tissue volumes corresponding to these images (see Supplementary Table 2) and assuming a typical cell is between 10 and 20 microns in width yields estimated numbers of cells per image of around 6,700 to 23,000, 1,900-15,000 and 2,900-53,000, in reasonable agreement with the number of final 3D cells. Extensive vascularization of thymus and lymph node may be expected to reduce the cell density.

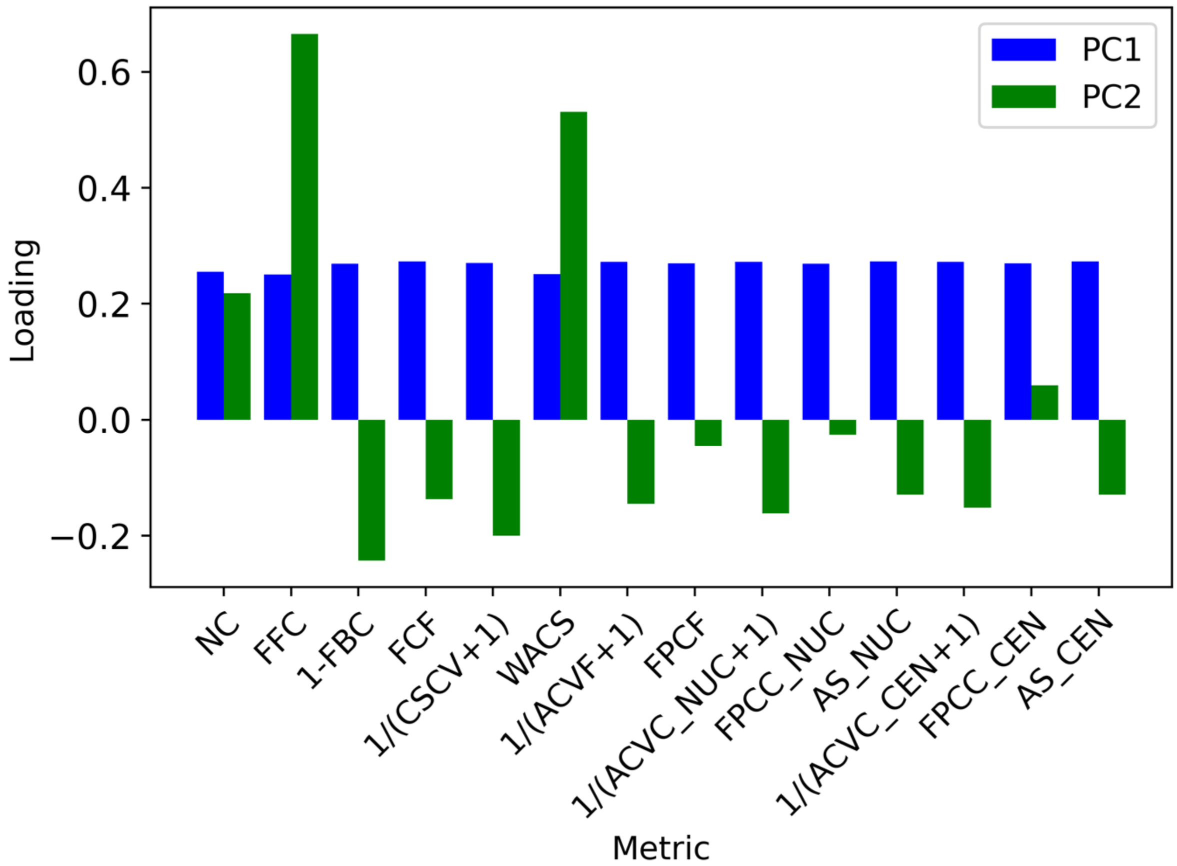


**Supplementary Figure 4** Factor loadings of PCA model trained by all 3D segmentation masks from all methods on hiPSC cell culture images.

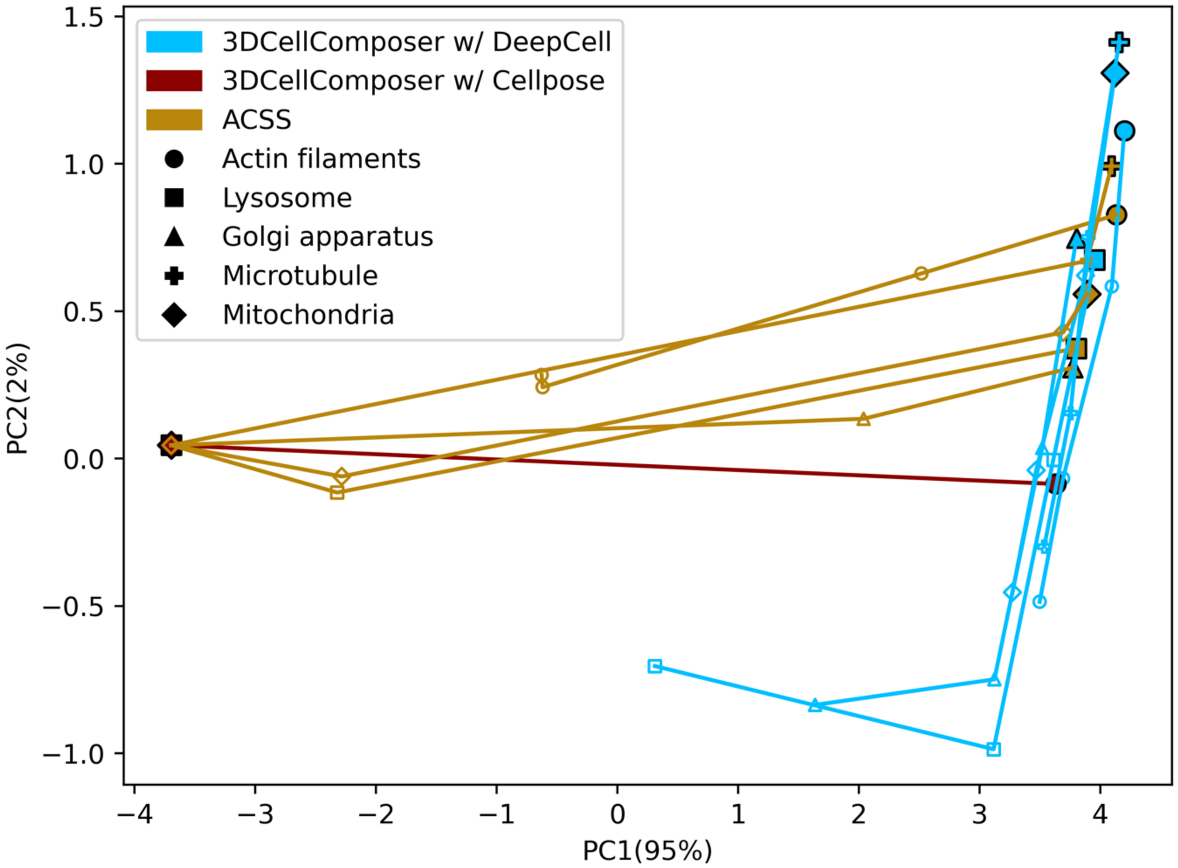


**Supplementary Figure 5** The top two principal components are illustrated for 3DCellComposer and ACSS on 3D hiPSC cell culture images for five cellular markers. Each method is denoted by a unique color, and distinct marker shapes correspond to the five cellular markers. Trajectories represent low, medium, high levels of zero-mean Gaussian noise perturbation on the images with standard deviations of 30, 60, and 90 voxel intensity, respectively. Here, a standard deviation of 90 corresponds to approximately 5% of the maximum intensity of the original images.

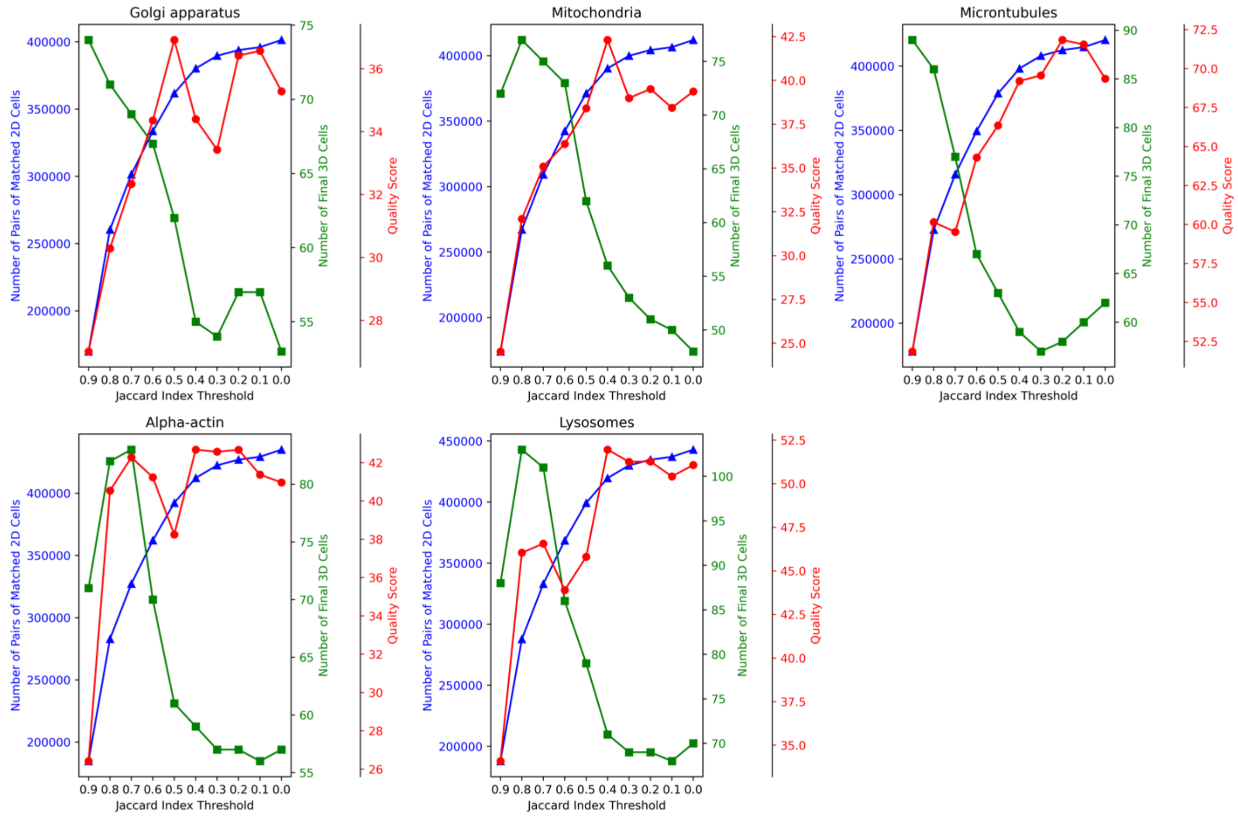


**Supplementary Figure 6** Analysis of the Jaccard Index optimization for 3DCellComposer using DeepCell with cell membrane input on hiPSC images. The field for these images is approximately 100 by 67 microns, or 6,725 square microns. Assuming the cells form a monolayer and a typical cell is roughly 10 by 10 microns gives an estimated number of cells per field of around 67, in good agreement with the number of final 3D cells for each dataset.
